# REAVER: Improved Analysis of High-resolution Vascular Network Images Revealed Through Round-robin Rankings of Accuracy and Precision

**DOI:** 10.1101/707570

**Authors:** Bruce A. Corliss, Richard Doty, Corbin Mathews, Paul A. Yates, Tingting Zhang, Shayn M. Peirce

## Abstract

Alterations in vascular networks, including angiogenesis and capillary regression, play key roles in disease, wound healing, and development. Imaging of microvascular networks can reveal their spatial structures, but effective study of network architecture requires methods to accurately quantify them using a variety of metrics. We present REAVER (Rapid Editable Analysis of Vessel Elements Routine), a freely available open source tool that researchers can use to analyze and quantify high resolution fluorescent images of blood vessel networks, and assess its performance compared to alternative state-of-the-art image analysis software programs. Top performing programs for each metric are identified by assigning a rank based on statistical multiple comparisons of accuracy and precision, modeled as matches in a round-robin style tournament. This comparison method yields a clearly defined and consistent standard for characterizing program performance, avoiding the use of non-standard ad hoc interpretations of multiple comparisons between programs. Using this comparison method and a dataset of manually analyzed images as a ground-truth, we show that REAVER was the top ranked program for both accuracy and precision for all metrics quantified, including vessel length density, vessel area fraction, mean vessel diameter, and branchpoint count. REAVER can be used to quantify differences in blood vessel architectures between study groups, which makes it particularly useful in experiments designed to evaluate the effects of different external perturbations (e.g. drugs or disease states).

## Introduction

The vast networks of interconnected blood vessels found in tissues throughout the body play significant roles in oxygen transport, nutrient delivery, and inflammatory response. The microvasculature is a key effector system in healthy and pathological conditions: serving primary roles in maintaining tissue homeostasis^1^ as well as the pathogenesis of disease^2^. The morphological structure of a microvessel network is closely intertwined with its biological functions, and quantitative changes in structure provide evidence of an altered physiological or pathological state. Examples include vessel diameter as an indicator of vasodilation, vasoconstriction, or arteriogenesis^2^, and vascular length density as an indicator of altered levels of tissue oxygenation^3^ or tissue regeneration^4^. Since the structural architecture of microvessel networks is closely intertwined with function, changes in microvessel architecture can, therefore, be used to assess cellular and tissue level responses to disease and treatments. Fluorescently labeled vascular networks can mark cells in a cell-type specific fashion at high signal-to-noise^5^, and when combined with widely available high-resolution imaging modalities such as confocal laser-scanning microscopy, serves as a well-established gold standard method for imaging cellular structures such as vascular networks^6^.

Several image processing programs have been previously used to quantify fluorescent images of microvessel architecture in an automated manner, including Angioquant^7^, Angiotool^8^, and RAVE^9^. While these programs have been used in various studies, they are estimated to have a low degree of adoption by the research community relative to the multitude of studies that have quantified microvascular architecture using a manual apprach^2^. Furthermore, the publications that introduce these tools for automation often provide nonstandard forms of metrics that make comparison between them difficult^2^ and lack a common method for evaluating performance. For image segmentation, manual analysis through visual inspection remains the gold standard technique^10–12^, defined as the method accepted to yield results closest to the true segmentation. Manual analysis can therefore be used as an approximation of ground-truth^13^ and a basis to compare performance of other automated analysis methods, with any perceived disagreement classified as error^14^.

In this paper, we establish and validate a new open source tool, named REAVER, for quantifying various aspects of structural architecture in fluorescent images of microvascular networks (Fig.1A) that uses simple image processing algorithms to automatically segment and quantify vascular networks while offering the option for manual user curation (Fig 1B). We use a benchmark dataset of fluorescently labeled images from a variety of tissues that exhibit a broad range of vascular architectures as a means of assessing each program’s general ability to automatically analyze vessel structure and minimize possibility of bias resulting from examining any single tissue. The error of REAVER’s output to ground-truth for various output metrics, including vessel length density, vessel area fraction, vessel tortuosity, and branchpoint count, is compared to other available vascular image analysis programs. Error is evaluated through accuracy of the output metrics, defined as the closeness of a measured value to ground-truth^15^, and measured with absolute error^16–18^. Precision, the random errors caused by statistical variability, is measured by comparing the variance of error between programs.

**Figure 1:**
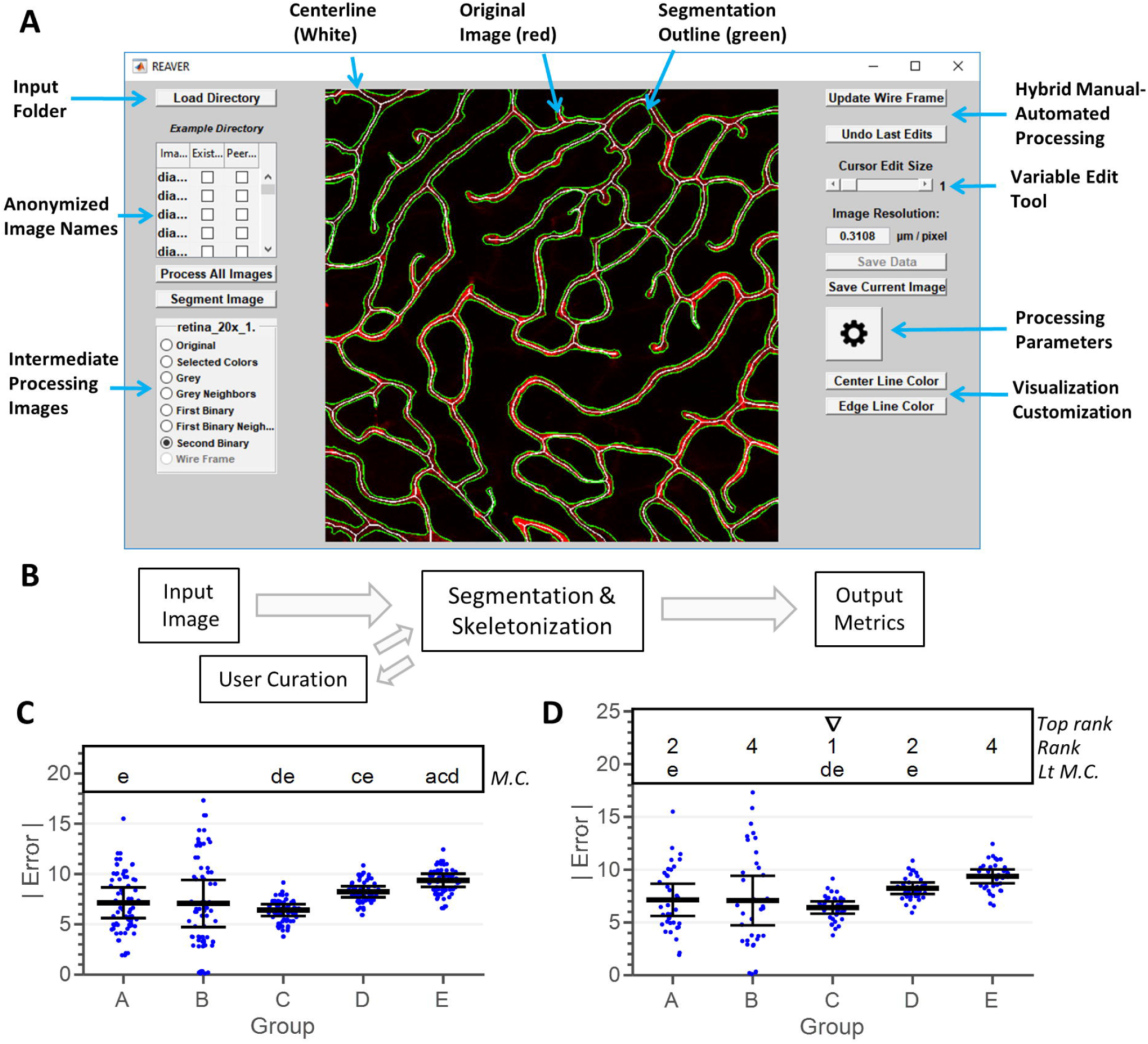
REAVER is an image analysis program for quantification of vascular networks in fluorescent images, while round-robin rankings allows for evaluating program error when multiple comparisons results are unclear. (**A**) Screenshot of REAVER graphical user interface. (**B**) Flow chart of data processing pipeline. (**C**) Comparison of absolute error between hypothetical image analysis programs, with letters denoting significant two-tailed relations in multiple comparisons (two-tailed pairwise t-tests, with Bonferroni correction of 10 comparisons, N=36 measurements, α=0.05, simulated data). (**D**) Identical dataset with proposed round-robin ranking analysis of performance (one-tailed less-than pairwise t-tests, with Bonferroni correction of 20 comparisons, N=36 measurements, α=0.05, simulated data). Annotations above graph: groups assigned top rank (triangle) when assigned the first rank (numbers) from a round robin tournament analysis of one-way less-than multiple comparisons (significance relations between groups denoted by letters).

An effective measurement technique exhibits high accuracy and precision with output metrics^15^, but traditional two-tailed multiple comparisons do not yield a clear interpretation of which programs have the lowest error among many since two-tailed tests yield non-directional conclusions^19^ and a lack of significant differences between programs can lead to ambiguous results. This is illustrated by comparing absolute error between hypothetical measurement methods A-E in Figure 3: there is a lack of standard interpretation of which groups are the best using post-hoc two-tailed multiple comparisons, especially with method A, B, and C that seem to have the lowest error with no significant difference between them (Fig. 1C). We developed a statistical method to identify the programs with lowest error by performing bi-directional one-tailed hypothesis tests, where we tested which groups had less error in a pairwise fashion, with comparisons serving as matches in a round-robin tournament between the programs. Comparisons with statistical significance were classified as wins, providing a basis to rank the programs based on their performance against other programs. In the example case, method C would be top ranked because it is significantly better than two other programs, both D and E, providing a standard interpretation of performance (Fig. 1D).

REAVER’s effectiveness is highlighted by being a top-ranking program across all metrics in both accuracy and precision. Given the ubiquity of high-resolution fluorescent microscopy and the established need for automated, rigorous, and unbiased methods to quantify vessel architectural features, we present REAVER as an image analysis tool to further microvascular research.

## Results

REAVER was developed to analyze and quantify fluorescent images of vessel architecture using basic image processing techniques, including adaptive thresholding and various filters for segmentation refinement (Supplementary Fifg. 1A-G, see Methods: REAVER Algorithm). A dataset of images was acquired from multiple mouse tissues (Supplementary Fig. 2A-F) and analyzed both manually and in an automated fashion using the REAVER, Angioquant^7^, Angiotool^8^, and RAVE^9^ software packages (Fig 2A-E). The metrics quantified from the manual segmentation, along with the segmentation itself, were used as ground-truth data. Any disagreement between the automated techniques to ground-truth was classified as error, allowing for comparison of performance between programs. Some of the programs had adjustable settings that altered the image analysis process: default image processing settings were used for all programs as a test of general performance with quantifying vascular architecture from fluorescently labeled images.

**Figure 2:**
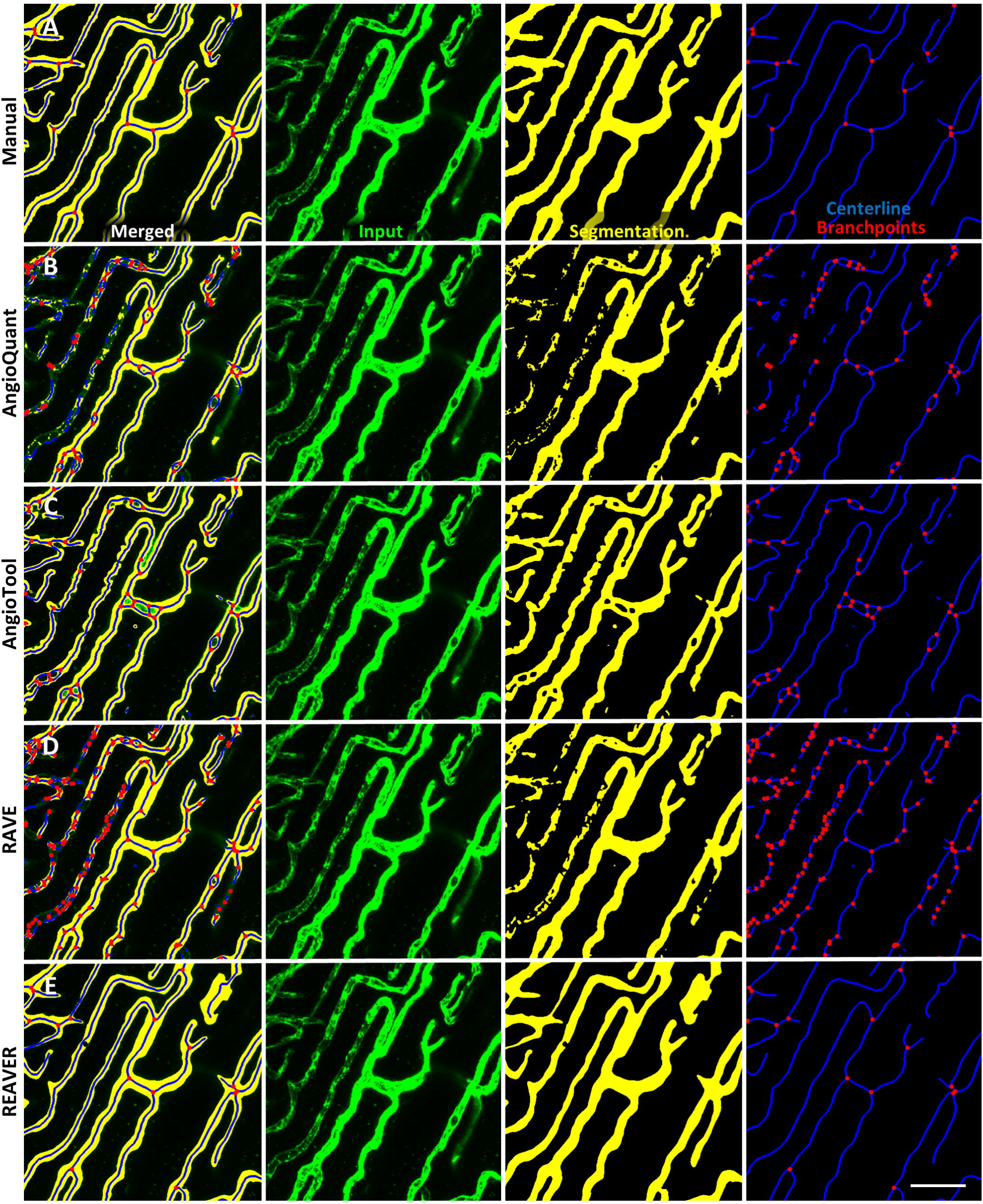
Manual segmentation and analysis can be used as ground-truth in comparing quantification pipelines. Representative image processed with (**A**) manual analysis, (**B**), AngioQuant, (**C**) AngioTool, (**D**) RAVE, and (**E**) REAVER with input (green), segmentation (yellow), centerline (blue), and branchpoints and endpoints (red) of vascular network (scale bar 50 um).

### REAVER demonstrates higher accuracy and precision across metrics

When the accuracy of vessel length density measurements was examined across the different automated image analysis tools (Fig. 3A), REAVER exclusively held the top rank, having a mean absolute error that was lower than all other programs (76.5% reduction with p=6.57e-3, AngioTool, one-tailed paired t-tests with Bonferroni adjustment). All programs except AngioQuant had evidence of a nonzero bias revealed through individual two-tailed t-tests for a mean of zero (p<0.05). When the precision of vessel length density measurements was compared across analysis programs, REAVER exclusively held top rank, having lower random error than all other programs (84.6% reduction with p=1.61e-3 from next closest group, AngioTool, one-tailed paired t-tests with Bonferroni adjustment) (Fig. 3B).

**Figure 3:**
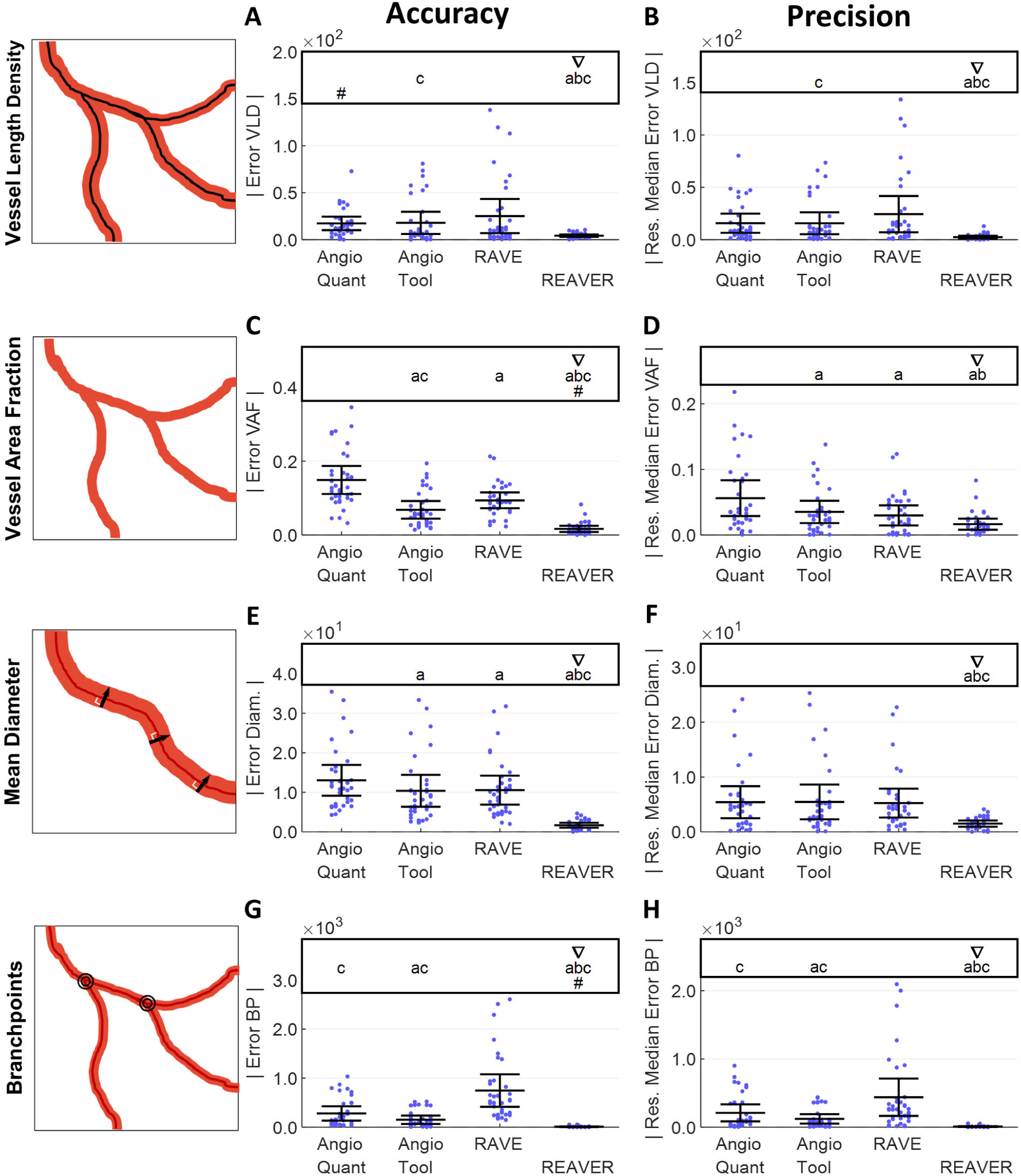
REAVER demonstrates top ranked accuracy and precision across metrics compared to alternative blood vessel image analysis programs. To evaluate accuracy, absolute error of (**A**) vessel length density (mm/mm^2^), (**C**) vessel area fraction, (**E**) vessel diameter (µm), and (**G**) branchpoint count compared to manual results (one-tailed less-than paired t-tests, forward and reverse order of all combinations with Bonferroni correction, 12 comparisons, α=0.05, N=36 images). For analysis of precision, the absolute value of residual error to group’s median error for (**B**) vessel length density (mm/mm^2^), (**D**) vessel area fraction, (**F**) vessel diameter (µm), and (**H**) branchpoint count (one-tailed less-than paired t-tests, forward and reverse order of all combinations with Bonferroni correction, 12 comparisons, α=0.05, N=36 images). For the annotations above each plot, top performers in each group defined by round-robin ranking of statistical outcomes (triangle), along with significant pairwise less-than comparisons between groups with Bonferroni adjusted p-values (letters). Groups are annotated when there is no evidence of nonzero bias with error, as determined by the origin falling within the bounds of the 95% confidence interval of the mean with Bonferroni adjustment of 4 comparisons (pound sign). Vessel metric diagrams were modified from^2^.

REAVER also held the top rank for accuracy with quantifying vessel area fraction, having a mean absolute error that was lower than all other programs (75.8% reduction with p=6.16e-8 from next ranked group, AngioTool, one-tailed paired t-tests with Bonferroni adjustment) (Fig. 3C). All programs except REAVER had a nonzero bias, revealed through individual two-sided t-tests for a mean of zero (p<0.05). When the precision of vessel area fraction was examined, REAVER was exclusively assigned the top rank, having lower random error than all other programs except RAVE (53.3% reduction with p=8.62e-3 from next ranked group with significant difference, AngioTool, one-tailed paired t-tests with Bonferroni adjustment) (Fig. 3D).

REAVER alone held the top rank with lower absolute error in vessel diameter than all of the other programs (83.9% reduction with p=8.29e-7 from next ranked group, AngioTool, one-tailed paired t-tests with Bonferroni adjustment) (Fig. 3E). All programs, including REAVER, exhibited evidence of nonzero bias revealed through individual two-tailed t-tests for a mean of zero (p<0.05). In terms of the precision of the vessel diameter measurement, REAVER was exclusively assigned the top rank, having lower random error than all other programs (72.3% reduction from next ranked group AngioQuant, with p=1.66e-3, one-tailed paired t-tests with Bonferroni adjustment) (Fig. 3F).

When it came to quantifying the accuracy of the branchpoint density measurement, REAVER was exclusively assigned the top rank, having a mean absolute error that was lower than all other programs (94.6% reduction with p=4.43e-5 from next ranked group, AngioTool, one-tailed paired t-tests with Bonferroni adjustment) (Fig 3G). All programs except REAVER had a nonzero bias, revealed through individual two-tailed t-tests for a mean of zero (p<0.05).REAVER was exclusively assigned the top rank for precision in quantifying branchpoint density, having lower random error than all other programs (93.2% reduction with p=4.70e-5 from next ranked group, AngioTool, one-tailed paired t-tests with Bonferroni adjustment) (Fig. 3H).

### REAVER exhibits higher segmentation accuracy and sensitivity with faster execution time

The error in the automated vessel segmentation was examined across all images in the benchmark dataset relative to the segmentation from manual analysis^20^. REAVER was exclusively assigned the top rank in segmentation accuracy, with a mean accuracy that was higher than all of the other programs (6.4% increase from next ranked group AngioTool,p=1.73e-7, one-tailed paired t-tests with Bonferroni adjustment) (Fig. 4A). In terms of sensitivity, REAVER was exclusively assigned the top rank with mean sensitivity that was greater than all of the other programs (34.1% increase from next ranked group AngioTool, with p=1.00e-15, one-tailed paired t-tests with Bonferroni adjustment) (Fig. 4B). In terms of specificity, RAVE and AngioQuant were assigned the top rank with a higher specificity than the other two programs (0.4% increase from next ranked group AngioTool, with p=4.39e-2, one-tailed paired t-tests with Bonferroni adjustment) (Fig. 4C). With regards to execution time, REAVER was exclusively assigned the top rank with a faster execution time compared to all of the other programs (36.4% reduction from next ranked group, with p=1.8e-16, one-tailed paired t-tests with Bonferroni adjustment) (Fig. 4D). All automated program execution times were <1% of the time required for manual analysis (3,089 ± 1,355 seconds per image, not displayed due to orders of magnitude difference), highlighting a major benefit of automated techniques.

**Figure 4:**
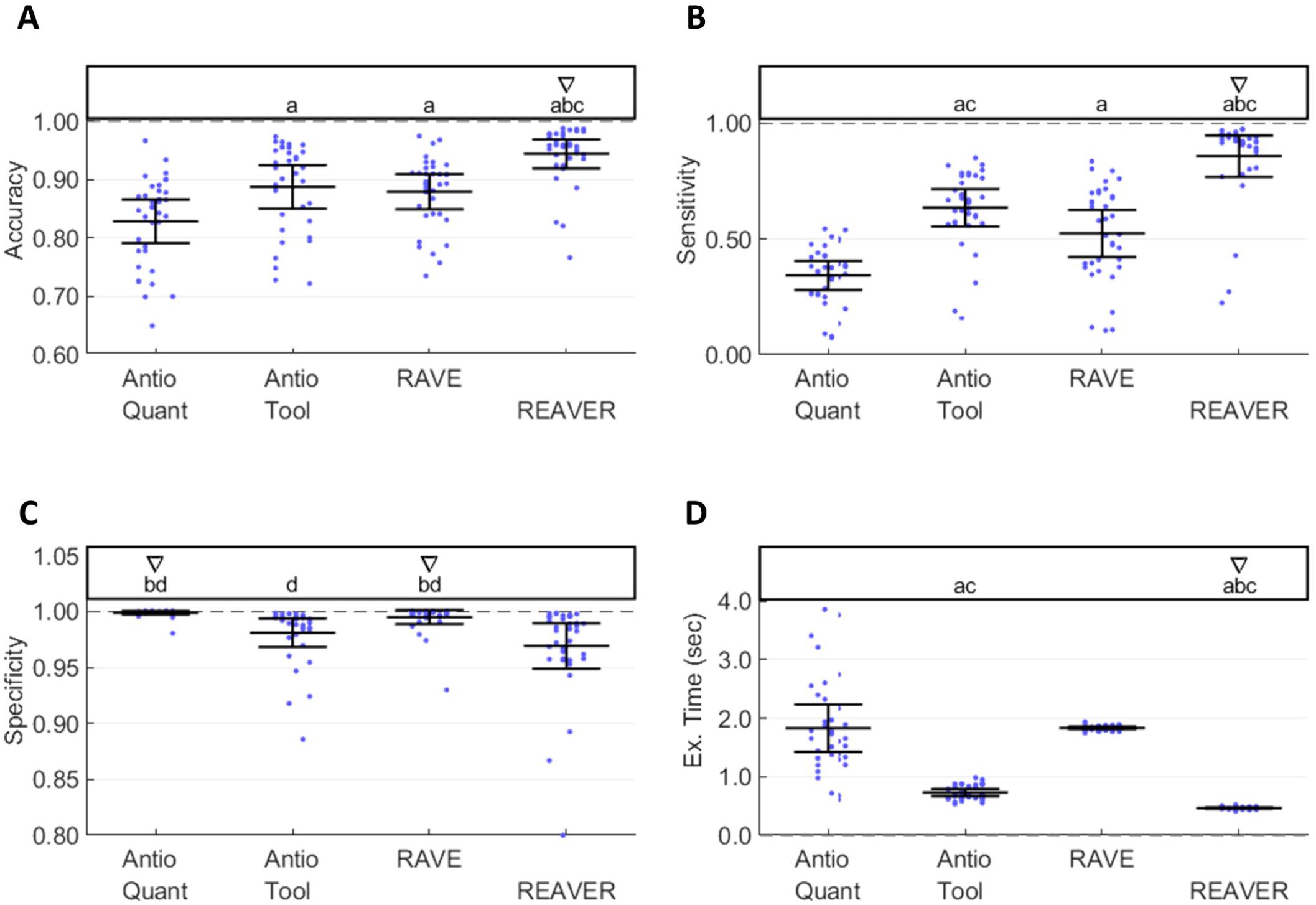
REAVER exhibits higher sensitivity and specificity with vessel segmentation, along with lower execution time compared to alternatives. Using the test dataset of images with manual analysis as ground-truth, the (**A**) accuracy, (**B**) sensitivity, and (**C**) specificity of the segmentation for each program, along with (**D**) execution time for each image (one-tailed greater-than (A-C) or less-than (D) paired t-tests, forward and reverse order of all combinations, with Bonferroni correction; 12 comparisons, α=0.05, N=36 images). For the annotations aboveeach plot, top performers in each group defined by round-robin ranking based on statistical comparisons (triangle), along with significant pairwise greater-than (A-C) or less-than (D) comparisons between groups with Bonferroni adjusted p-values below significance level (letters).

### Blinded manual segmentation curation can improve accuracy of metrics

The errors for each of the output metrics relative to the manual analysis were compared for: 1) metrics obtained by REAVER using purely automated analysis, and 2) metrics obtained by using a combination of automation paired with manual curation of the image segmentation. Using the same images and internal image processing parameters (as used in Figure 2), the absolute error across all images were compared before and after manual curation where the user was blinded to the group each image belonged to. The absolute error for vessel length density was reduced 45% (p=6.4E-5, paired two-tailed t-test with Bonferroni adjustment, Fig. 5A) while there was no change to the vessel area fraction error (p=1, paired two-tailed t-test with Bonferroni adjustment, Fig. 5B). Absolute error in vessel diameter measurements had a decreasing trend, with a 25.0% reduction in absolute error (p=0.188, paired two-tailed t-test with Bonferroni adjustment, Fig.5C), and absolute error in branchpoint density measurements experienced a similar decreasing trend with 17.7% reduction in absolute error (p=0.112, paired two-tailed t-test with Bonferroni adjustment, Fig. 5D).

**Figure 5:**
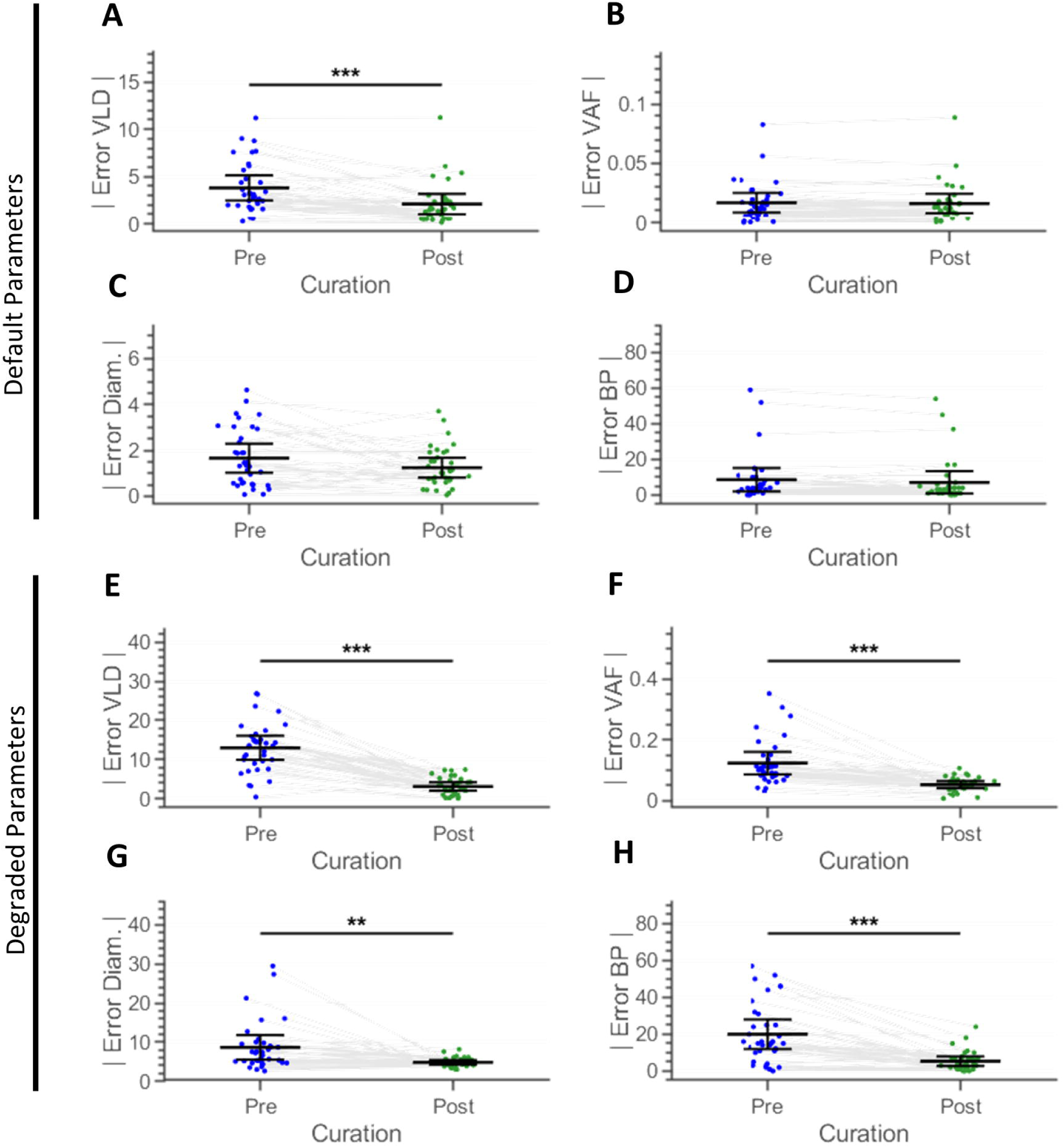
Image segmentation curation can enhance accuracy of output metrics. Comparison of error with (**A**) vessel length density (mm/mm^2^), (**C**) vessel area fraction, (**E**) vessel diameter (µm), and (**D**) branchpoint count from automated analysis using default parameters before and after manual curation of image segmentation. Comparison of error with (**E**) vessel length density, (**F**) vessel area fraction, (**G**) vessel diameter, and (**H**) branchpoint count from automated analysis using degraded parameters before and after manual curation of image segmentation (for each of the two datasets, two-tailed paired t-tests with Bonferroni correction, 4 comparisons, α=0.05, N=36 images).

Since REAVER demonstrated superior performance with this image dataset compared to the other programs, the error for many of the metrics was small, consequently lowering the potential effect size that manual curation may provide. To test if manual curation is useful for lower quality results that could benefit more from manual curation, REAVER’s internal image processing parameters were intentionally set to extreme values to produce a heavily flawed segmentation. Using the same dataset of images, user curation increased the accuracy for all of the metrics: the absolute error for vessel length density was reduced by 75.9% (p=1.64e-11, pared two-tailed t-test with Bonferroni adjustment, Fig. 5E), the vessel area fraction absolute error was reduced 57.5% (p=9.99e-6, pared two-tailed t-test with Bonferroni adjustment, Fig. 5F), vessel diameter absolute error was reduced 44.5% (p=4.79e-3, pared two-tailed t-test with Bonferroni adjustment, Fig. 5G) and branchpoints absolute error was reduced by 73.2% (p=1.36e-6, pared two-tailed t-test with Bonferroni adjustment, Fig. 5H).

### REAVER reveals differences in microvascular architectures across spatial locations in murine retina

An effective microvascular image analysis program can separate between groups of images with known differences in microvascular architecture. The blood vessels of the murine retina are a well characterized microvascular network that exhibits extensive heterogeneity of vessel architecture depending location in the tissue^2^, both with radial distance from optic disk, and with each of the three discrete layers of vasculature beds: the deep plexus, intermediate plexus, and superficial capillary plexus^21^. With a dataset of images separated by two radial distances from the center of the retina, at each of the three vascular layers (Supplementary Figure 3A, B), REAVER could discern unique vessel architectural features across the metrics quantified (Supplementary Figure 3D-L). These metrics were able to achieve a partial linear separation between retina locations with the first two components of a principle components analysis (Supplementary Figure 3C).

## Discussion

We present a novel software package, REAVER, for quantifying metrics that have been classically used to describe microvascular network architectures. Through a novel application of round-robin ranking, we simultaneously examined and ranked the accuracy and precision of REAVER compared to three published automated image analysis tools. This comparison revealed that REAVER is the top ranked program in terms of accuracy and precision across all four metrics of vessel architecture examined. We believe this is explained by the fact previous programs were originally developed when basic image processing algorithms were computationally expensive^8,9^, or for other image modalities (such as vascular images from transmission light microscopy^7^). REAVER may have outperformed the other programs because of its higher degree of accuracy with automated image segmentation, yielding a segmented structure closer to truth than the other programs. REAVER’s higher sensitivity in segmentation demonstrated in this dataset highlights its improved ability to correctly discern foreground pixels of vessel architecture, while it’s lower performance in specificity suggests that other programs are better at correctly discerning background non-vessel pixels. For this particular application, we argue that sensitivity and specificity are not as important as accuracy for evaluating a program’s overall performance: higher sensitivity can simply be accomplished with over-segmenting the vessel architecture, while higher specificity can be attained by under-segmenting the image. Although REAVER had the fastest execution time compared to other programs, we argue that all programs demonstrated acceptable execution times given the low cost of computational processing power^22,23^.

Performance of image analysis programs can be examined with the Bland-Altman analysis (Supplementary Fig. 4 A-P), a technique that compares two measurement methods based on paired measures^24,25^ and establishes agreement if the range of the agreement interval (encompassed by the inner 95% span of the distribution of error between the two techniques) has an acceptable magnitude based on a priori defined limits that are application-specific^26^. This analysis is often used to compare a new measurement method with a previously developed gold standard method in order to test whether the new technique can be used in place of the previous one. Although some studies assert Bland-Altman is the only correct technique to compare methods of measurement^27^, we highlight that it does not provide a means to compare performance of multiple measurement methods to ground-truth. The 16 Bland-Altman plots generated across the 4 programs and 4 metrics tested yields little insight into how well these measurement techniques performed relative to one another. Furthermore, it is frequently left up to the user to define the acceptable range for the agreement interval: in the absence of standardized approaches, this process can be influenced by perspective, opinion, and bias.

While our results present a relatively straightforward case where one program performed better relative to the others in terms of ranking for accuracy and precision, our analysis approach can be useful in investigations where the results across metrics are less consistent. Since multiple programs can be assigned top rank, this technique can be used as image analysis pipelines mature and the remaining effect size for potential improvement to ground-truth diminishes. We note that while the top ranked programs are the top performers, there are cases where it could be argued that lower ranked programs should be considered as well, such as in Fig. 4D where RAVE exhibited better precision than one of the programs and was not discernably different than REAVER, but REAVER alone held top rank because it had better precision than two of the other programs. We recommend that instances such as these be examined on a case-by-case basis that takes into account effect size and potentially adjusting the significance level for the ranking comparisons. Regardless, by identifying the best programs for accuracy and precision for each metric, researchers can prioritize program performance based on the most relevant metric for a specific biological hypothesis. For example, if a hypothesis is focused on changes to blood vessel density that is observed in pathologies like diabetes^28^, then performance of the program in terms of vessel length density would be prioritized in selecting the best program for this application. For studies focusing on developing better general tools to quantify vascular architecture, this framework will allow for comparing the performance of new software packages to preexisting versions, and provides a means to select for programs that have acceptable performance, defined as being a member of the top rank. A program within the top ranked group would provide justification for use, or for a more focused and detailed post-hoc paired examination using techniques like Bland-Altman analysis to examine effects of systemic and proportional error in more detail.

While automated results have the benefit of minimizing human interaction time with processing images and maintaining an unbiased analysis of data, there are instances where image segmentation may perform poorly and a higher degree of accuracy is needed. To accomplish this, we propose that manual curation of segmentations derived from automated analysis, with the user blinded to group assignment, would reduce error of output metrics. While manual curation of images using default image processing parameters for REAVER showed little improvement in accuracy across all metrics, the potential effect size for improvement was small due to REAVER’s high level of accuracy and precision. To probe whether manual curation can enhance quality of results, the same images were processed using extreme values for image processing parameters that led to a poor segmentation. Manual curation reduced mean absolute error approximately 60% across metrics, demonstrating the utility of hybrid approaches of data analysis where automated and manual techniques can be combined to enhance data quality. Our results indicate there are cases where manual curation can range from showing little benefit in enhancing data quality to profoundly improving the accuracy and precision of results. Using a pilot study of a small dataset of images comparing both automated results and automation with curation to ground-truth will reveal to a researcher if curation is worth the time investment for a particular application. Furthermore, manual curation of automated segmentation represents a promising technique for efficiently generating ground-truth analysis of images that requires much less time than purely manual techniques. It is important to note that we only investigate each program’s ability to automatically segment the vasculature: many of them include several manually adjustable image processing settings (although none offer the option for direct manual curation), and there is a possibility that one of the other programs would perform better than REAVER with optimal parameters. Testing performance with manual adjustments would be a complex undertaking reserved for future research, requiring not only a fair method for identifying optimal parameters for each image and program under realistic use cases, but also evaluating how effective a user can be at identifying the optimal parameters and obtaining the optimal segmentation.

While our comparison of the precision and accuracy of four different automated image analysis programs was achieved by performing a separate comparison for each metric, in the future, it would be beneficial to compare program performance across all metrics simultaneously. This could require a method of weighting metrics based on a metric’s ability to discern alterations in a relevant biological dataset, while accounting for covariance and dependence between metrics (such as vessel length density being closely correlated with vessel area fraction for vessel networks with nearly uniform vessel diameters). Program performance could also be compared by combining the examinations of accuracy and precision for all metrics into a multi-round-robin tournament and calculating rank across them^29^. The evaluation of trueness or bias, defined as the average distance between an output metric across images and ground-truth values^15^, is not included in this study because no method exists to statistically compare trueness between study groups since distributions must be compared to each other and their distance to zero bidirectionally at the same time. The development of such a technique would be required for discerning differences in trueness and lead to a more complete characterization of error and performance of the programs examined. Furthermore, our representation of ground-truth could be improved by having multiple users manually analyze the images to generate a gold standard from the consensus, as done previously with image object classification^30^.

In summary, we introduce REAVER, a new software tool for analyzing architectural features in microvascular networks, that achieved top ranked accuracy and precision for all metrics quantified in our study. We introduce a method for quantitatively comparing accuracy and precision using one way pairwise comparisons analyzed with round-robin tournament ranking to identify top performers. Both of these contributions fill an existing void by providing a better image processing program for blood vessel architecture quantification, as well as a framework for evaluating effectiveness and performance of measurement techniques in an unbiased manner.

## Methods

### Code and Data Availability

REAVER Source code available under a BSD 3.0 open source license at: https://github.com/bacorliss/public_REAVER. It was written MATLAB 2018a and requires the image processing toolbox to run. Benchmark image dataset and annotations are publicly available and linked from the repository main page.

### Murine Retinal Harvest

All procedures were approved by the Institutional Animal Care and Use Committee at the University of Virginia, and completed in accordance with our approved protocol under these guidelines and regulations. We used C57Bl6/J mice from The Jackson Laboratory (JAX stock #000664, Bar Harbor, ME). Mice were sacrificed via CO2 asphyxiation with cervical dislocation for secondary sacrifice, eyes enucleated, and incubated in 4% PFA for 10 minutes. Single mice were used for the 36-image dataset across tissues (Fig. 3-4, Supplementary Fig.1) and for the retinal location dataset (Supplementary Fig. 3) to examine vascular heterogeneity within a single biological sample.

### Immunohistochemistry and Confocal Imaging of Retinas

Retinas were labeled with IB4 Lectin Alexa Flour 647 (ThermoFisher Scientific, Waltham, MA, I32450) and imaged at using a 20x objective (530 um field of view) and a 60x objective (212 um field of view) with a Nikon TE-2000E point scanning confocal microscope. A total of 36 z-stacks from six different tissues were flattened to 2D images with a maximum intensity projection and used as a benchmark dataset for segmenting vessels and quantifying metrics of vessel architecture. To establish ground-truth, all images were manually analyzed in ImageJ.

### REAVER Algorithm

REAVER’s algorithm was implemented in MATLAB and designed to process the image in two separate stages: segmentation based on intensity over local background and then skeletonization and refinement. Segmented vasculature is identified through a combination of filtering, thresholding, and binary morphological operations. The image is first blurred with a light blurring averaging filter with an 8-pixel neighborhood, and then an image of the background (low frequency features) is calculated with a larger user-defined heavier averaging filter (default: 128 pixels). To create a background-subtracted image, the heavily blurred background image is subtracted from the lightly blurred image. The background-subtracted image is thresholded by a user-defined scalar (default: 0.045) to generate an initial segmentation. Next, the segmentation border is smoothed and extraneous pixels are removed with an 8-neighborhood convolution filter that is thresholded such that only pixels with at least 4 neighbors are kept. Leveraging the domain-specific knowledge that vessel networks are comprised of large connected components, those with area less than a user-defined value are removed (default: 1600 pixels). To further smooth segmentation borders, the complement of the segmented image is convolved with an 11-square averaging filter and values are thresholded above 0.5. To fill in holes within segmented vasculature, connected components of the complemented segmentation with less than 800 pixels are set to true. The images are then thinned to compensate for a net dilation of segmentation from earlier processing steps. Finally, connected components of size less than a user-set value (default: 1600 pixels) are removed again to generate the final segmented image.

To generate the vessel centerline, the segmented image border is further smoothed with eight iterative applications of a 3-pixel square true convolution kernel thresholded such that pixels with at least 4 neighbors are set to true. To fill in small holes and further clean the segmentation edge, the MATLAB binary morphological operations “bridge” and “fill” are applied in that order four times, along with an application of a 3-pixel square majority filter where every pixel needs 5 or more true pixels in the square to pass. Connected components in the complement of the segmentation with pixel area less than 80 pixels are set to true in order to fill in holes within segmented vessels. The initial vessel centerline is identified by applying the binary morphological ‘thin’ an infinite number of times to the segmentation with replication padding applied; otherwise thinned centerlines would not extend to the end of the image.

To filter out centerlines for segments that are too thin, a Euclidean distance transform is calculated from the complement of the segmented image and sampled at the pixel locations of the vessel centerline, resulting in a thickness centerline image where the vessel centerline contains values for the radius of the vessel in that region. The thickness centerline is divided into individual vessel segments via its branchpoints, the average radius calculated for each, and segments that fall below the user-defined thickness threshold were removed (default: 3). The refined vessel centerline was further cleaned with MATLAB’s “spur” and “clean” morphological operations, along with a final morphological thinning. Branchpoints and endpoints are identified with MATLAB’s built in morphological operations, ignoring features located at the image border because edge effects cause false positives.

We note that while this algorithm was tested with a benchmark image dataset that included a practical range of resolutions with the default image processing set of parameters, the parameters are resolution-dependent to some degree. We argue that the resolution range we used, images acquired at 20x and 60x magnification, represent the most relevant range of modalities for probing complete microvascular structures. Lower magnification below 20x lacked sufficient resolution to discern the structure of the smallest vessels of the microvascular network, while higher magnification over 60x sampled such small areas of vasculature that estimates of various metrics of vascular structure would be unreliable. Using resolutions far outside this range would require changing the default image processing parameters.

### Manual Analysis of Benchmark Dataset

To make the time demands for establishing ground-truth manageable, a mixed-manual analysis approach was used to analyze the benchmark dataset, where a simple set of ImageJ macros provided an initial guess for thresholding and segmenting blood vessels, and then the user manually used the paintbrush to draw in changes required. The initial automated guess was used to save time, but there is a possibility that it biased the ground-truth data to unfairly favor REAVER’s results. To check if bias in ground-truth could alter statistical outcomes, a completely manual segmentation was compared to the mixed manual method in a subset of images from the benchmark dataset (N=6 images, one from each tissue type, Supplementary Fig. 5A-D). The completely manual analysis was conducted by a different user with no cross-training between those who did the mixed manual analysis to represent the worst-case estimation of disagreement between the two methods. The disagreement of four output metrics (vessel length density, vessel area fraction, vessel diameter, and branchpoints) was examined via Bland-Altman plots, and all metrics had no evidence of bias (N=6 images, p-values displayed in each chart, Supplementary Fig. 5E-H, no Bonferroni correction applied for conservative interpretation). The width of the confidence intervals of the mean was calculated based on the 6 sample images (normality approximation, Supplementary Fig. 5, ObsW CI_95_). Since the confidence interval is based on the standard error (and decreases by 1/√n), the confidence intervals for the entire benchmark dataset is estimated based on increasing the sample size from 6 to 36 images with sample standard deviation fixed (Supplementary Fig. 5, EstW CI_95_). We found these estimated confidence intervals were minor in size compared to the effect sizes observed with the mean absolute error of the automated segmentation between the programs tested (Supplementary Fig. 5, columns labeled AngioQuant - REAVER).

The mixed manual analysis used for ground-truth for the benchmark dataset was acquired through manual curation of an initial automated threshold using macros in ImageJ to provide an initial guess of what structures in the image were considered vessels. Each image was loaded into ImageJ and an initial segmentation was calculated as a basis for manual curation. The image was segmented using a macro that removed high frequency features, applied local thresholding using the Phansalkar method^31^, decreased noise with the despeckle function, removed binary objects of pixel area less than 100 pixels, morphologically opened the image (erosion followed by dilation), applied a median filter on the adjacent four pixel neighborhood, and finally enhanced the brightness of the image for visibility.

Following this initial segmentation, trained editors used the paintbrush tool to correct errors in the segmentation. The total time to correct the segmentation was recorded. After the segmentation was adjusted to satisfaction, another ImageJ macro was run to generate a preliminary skeleton of the image. This script applied a median filter of radius 9 and the ImageJ Skeletonize operation. Once again, the curator used the paintbrush tool to correct the automatically generated skeleton. Special care was taken to ensure the skeleton had a width of only one pixel. The total time to correct the skeleton was recorded. The segmentation was run through the same analysis code that the other automated methods were analyzed with. The curator then tagged each branchpoint in the skeleton and recorded the total count and locations. This data was used as ground-truth to compare the automated analysis of several vessel architecture image processing pipelines.

### Image Quantification of Benchmark Dataset

Each software package provided different collections of metrics calculated in different ways. To fairly evaluate program performance in an unbiased fashion, a collection of four metrics was selected that could be calculated from the output data supplied by each program: specifically, the segmented vasculature image and the vessel centerline image. These output images were collected from each program, and then analyzed with the same code to quantify the vessel length density, vessel area fraction, mean vessel diameter, and number of branchpoints. If these output images were not available in the program, we either inserted code to export them to disk, or capture them from the program graphical display.

Angiotool is an open-source package written in JAVA. We could not successfully recompile the program to access and export the output images directly, and therefore had to use indirect means to obtain output images. An image was imported and processed with default settings (vessel diameter: 20, vessel intensity: [15,255], no removal of small particles, and no filling of holes). Images of the segmentation and centerline were derived by adjusting the display settings of the post-processed image and exported using the built-in Windows Print Screen function to capture images without distortion or compression. AngioQuant is written in MATLAB, and the source code was modified to output the segmented image and vessel centerline. Input images were inverted prior to importation to AngioQuant, and the default batch image processing parameters (kernel size: 1, edge tubules were not removed, and prune size: 10). All other numeric values were the default values for the program. RAVE was written in MATLAB, and the source code was modified to directly export the generated segmentation and skeleton for quantification. Each image was processed individually with default settings.

REAVER was written in MATLAB, and outputs the segmentation vessel centerline images in datafiles that are stored in the same directory as the analyzed images. All images were processed in batch mode with the default values (Averaging Filter Size: 128, Grey to Binary Threshold: 0.045 Minimum Connected Component Area: 1600, Wire Dilation Threshold: 0, and Vessel Thickness Threshold: 3). Once all images were processed and the associated mat files created by REAVER, the output images were extracted from these datafiles.

Once all 36 composite images containing the segmented image and the image centerline for each program were generated, a MATLAB script was used to calculate the values for the metrics from the composites. The vessel area fraction was calculated as the fraction of true pixels in the segmentation image. Vessel length density was calculated first by obtaining vessel pixel length through summing up all pixels in the vessel centerline images, converting this to millimeter units using the image resolution, and then dividing this by the image field of view in units of mm^2^. To calculate mean vessel diameter, a Euclidean distance transform of the segmentation channel was calculated where each pixel’s value was equal to its distance from the nearest false or un-segmented pixel. Then, the skeleton channel was used as a mask to sample the distance values corresponding to the vessel centerlines to obtain the radius of vessel segments. These values were multiplied by two and subtracted by 1 to get diameter values, and were converted to micrometer lengths using the image resolution. The MATLAB binary branchpoints morphological operation was used to find the branchpoints, and the number of branchpoints was calculated.

For analyzing the performance of the segmentation, true positive (TP) values for the image segmentation were calculated by taking the sum of the number of pixels that the program marked as true and that were also marked as true in the manual segmentation. True negatives (TN), false positives (FP), and false negatives (FN) were calculated and used to measure segmentation accuracy, sensitivity, and specificity (See Methods: Program Evaluation Metrics).

### Image Processing Execution Time

The processing times for the manual data were recorded using a stopwatch while the curator was editing the segmentation and skeleton images in ImageJ. The processing times for AngioQuant, RAVE, and REAVER were all collected by adding tic/toc statements that logs execution time into their MATLAB codes immediately before processing began and immediately after processing finished. This generated measurements for each program which were recorded.

Since AngioTool was provided as an executable file and the source code could not be successfully compiled without editing the code for dependency issues, reorganizing the file structure, and downloading external required libraries, the processing times were collected differently than the other three programs. The third-party application “Auto Screen Capture” (https://sourceforge.net/p/autoscreen/wiki/Home/) was used to capture images of the AngioTool application’s progress bar approximately every 15ms starting from before the start of processing to after it finished. The screenshots were automatically named as the exact time they were taken at the resolution of 1 ms. The collection of screenshots was inspected to identify the start time for processing based on the mean time of the final screenshot before the progress bar changed and the one immediately after. The end time for processing was determined by taking the mean time between the final image before the progress bar completed and the image immediately after. The difference between these two mean times was taken to get a total processing time. The total measurement error from collecting processing times in this way works out to be less than 3% of the total processing time.

All processing times were gathered on a computer with 32GB of DDR4-2666 RAM with CAS Latency of 15, an Intel i7-8700K 3.7 GHz 6-Core Processor, and a GeForce GTX 1080 graphics card with 8GB of VRAM. No overclocking, parallel processing, or GPU processing was used.

### REAVER Curation Analysis

REAVER’s code was modified to include a timer object which triggered every 20 seconds to save data to disk in the same manner as when manually specified. This timer started as soon as the curator used REAVER’s automatic segmentation and finished when the curator saved the curation results. After the automatic segmentation finished, the curator manually edited the segmentation using REAVER’s GUI and periodically updated the wire frame button. The accuracy of metrics from the pre-curation output were compared to post-curation output with output from the manual analysis serving as ground-truth. This process was initially conducted with default parameter values for REAVER’s image processing, but this yielded error with very small effect size across metrics, making it difficult to test for the potential benefits of manually curating automated results.

To test if image curation could help with lower quality image analysis results, this process was repeated with extreme shifts in default parameters, leading to a highly sub-optimal set of parameters that artificially created a lower quality segmentation with larger effect size for error to ground-truth (Averaging Filter Size: 64, Grey to Binary Threshold: 0.07). Additionally, within the image segmentation algorithm, reducing the extent of background subtraction, and the smoothing filter was changed to a minimum of 6 neighbors instead a minimum of 4 to yield a true pixel.

### Program Evaluation Metrics

The accuracy of the vessel structure metrics, defined as the closeness of a measured value to a ground-truth^15^, was examined with absolute error^16–18^ (Fig 3 A, C, E, G). Let *Y_i,j_* be the value of a given vessel structure metric (vessel length density, vessel area fraction, branchpoint count, and vessel diameter) from the *i^th^* image and *j^th^* program, and *G_i,j_* be the corresponding ground-truth value derived from manual analysis. We define error, *E_i,j_*, as the difference between a measurement and its corresponding ground-truth, and assess accuracy with the absolute error, *Ai,j*:

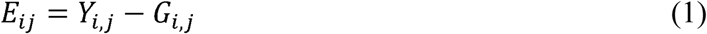

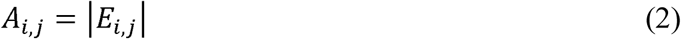

Measurements with low absolute error are considered highly accurate. We define precision^32^ *P_i,j_* of the *j^th^* program for *i^th^* image to be

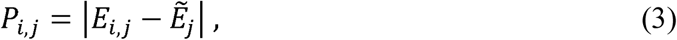

where 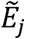 is the median of *E_ij_* across images, with *i*=1, …, 36 images, using the variable transform from the Brown-Forsythe test of variance^32^ (Fig 3 B, D, F, H).

Additionally, we evaluated metrics that quantify the agreement between each program’s vessel segmentation and the ground-truth (i.e., manual segmentation) across the entire image, evaluating accuracy (S^A^), specificity (S^C^), and sensitivity (S^N^) (Figure 4A-C). The definitions of these metrics depend on four quantities: true positives (TF), defined as the number of pixels classified as vasculature that agree with manual segmentation, true negatives (TN), the number of pixels classified as background in agreement with the manual segmentation, false positive (FP), the number of vasculature pixels in disagreement with manual analysis, and false negative (FN), the number of background pixels in disagreement with manual analysis. The metrics are^33^:

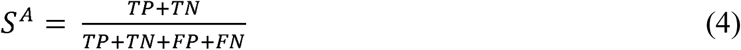

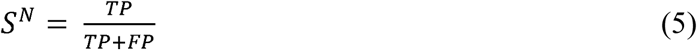

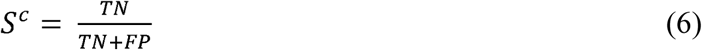

For evaluating the effectiveness of manual user curation of automated segmentation, we compared the accuracy before and following user curation of automatically processed images for each vessel structure metric (vessel length density, vessel area fraction, branchpoint count, and vessel diameter) with REAVER (Fig. 5A-D). Let *Y^B,D^_i,r_* denote the value of a given vessel structure metric before any user curation (superscript *B*) using default image processing parameters (superscript *D*) for the *i^th^* image from REAVER (program index *j* program set to *r*, the index for REAVER), and *G_i,r_* be the corresponding ground-truth value (as defined previously). The absolute error *A^B,D^_i,r_* used to evaluate accuracy would be defined as

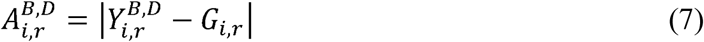

Let *Y^F,D^_i,r_* denote the value of a given vessel structure metric following user curation (superscript *F*) using default image processing parameters (superscript *D*) from REAVER (program index *j* set to *r*, the index for REAVER). The absolute error *A^F,D^_i,r_* is

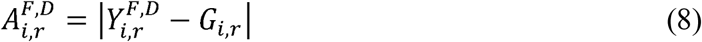

Error was also examined before and after user curation with a different set of internal image processing parameters set to substandard values (Fig. 5E-H). Let Y*^B,S^_i,r_* denote the value of a given vessel structure metric before any user curation (superscript *B*) using substandard internal image processing parameters (superscript *S*) from REAVER (program index *j* set to *r*, the index for REAVER). The absolute error *A^B,S^_i,r_* is

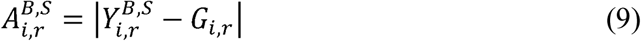

Let *Y^F,S^_i,r_* denote the value of a given vessel structure metric following user curation (superscript *F*) using default image processing parameters (superscript *S*) from REAVER and not any other program (with *j* set to *r*, the program index for REAVER), The absolute error *A^F,S^_i,r_* is

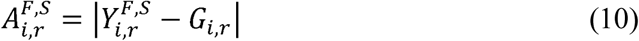

### Summary of Metric Classes

Metrics used in this study are split into two main classes (Table 1). Vessel structure metrics are the measures that describe architectural features of a vessel network and used for biological research. Program evaluation metrics are measures of error calculated from vessel structure metrics or derived from differences between each program’s image segmentation and corresponding manually segmented image. Program evaluation metrics are specifically used to compare error between programs and determine performance. To clarify the notation used for examining error before and after manual user curation, conventions are illustrated in Table 2.

**Table 1:**
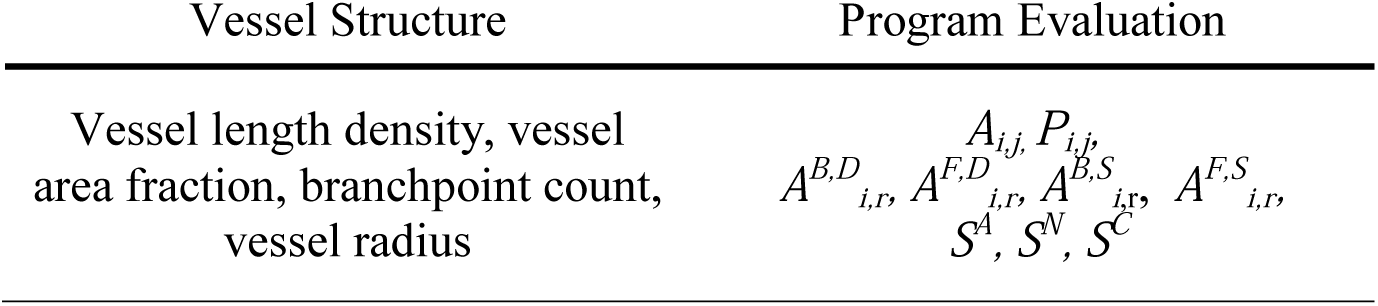
Metric Classes

**Table 2:**
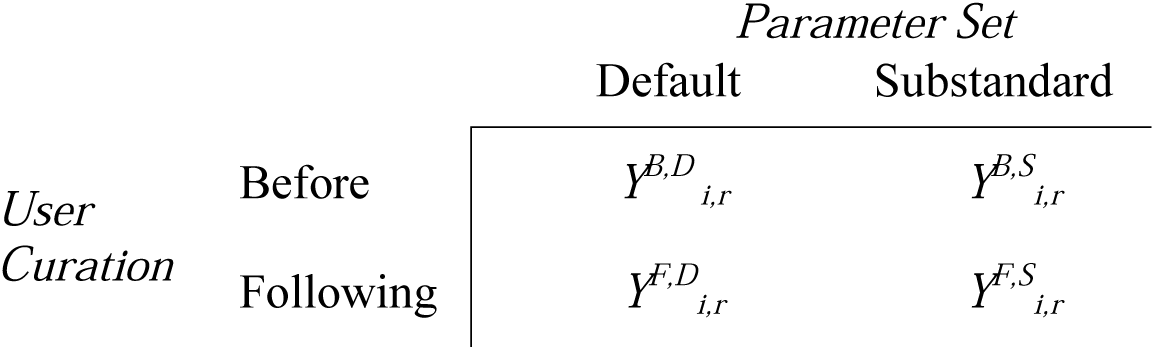
Metrics for Examining User Curation

### Statistical Analysis

To probe how programs performed relative to one another, we compared the distributions of absolute error *A_i,j_* and precision *P_i,j_* for all pairs and ordering of programs via one-sided paired t-tests (Fig 3). Specifically, for every two programs I and II, we tested both whether program I is better than program II and whether program II is better than program I. One-sided tests were used to determine which program among the two was better: that is, programs with vessel structure metrics exhibiting lower mean absolute error, standard deviation of error (Fig. 3), and execution time (Fig 4D) were preferable, while programs with higher segmentation accuracy, specificity, and sensitivity (Fig. 4A-C) considered better. We categorized these comparisons as wins and losses in a round-robin style tournament, assembled them into a dominance matrix, and ordered them through previously developed ranking algorithms^34^ to determine the best programs. (see Methods: Ranking Algorithm for detailed discussion). Members of the top rank for a given metric were annotated with a triangle above the plot.

For the 4 programs under study, we performed 12 one-sided tests in total simultaneously to compare them in a pairwise fashion with ordering in both directions. P-values were corrected with a Bonferroni adjustment^35^ using the following equation:

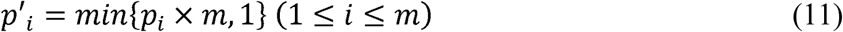

Where *p_i_* is the unadjusted p value, *m* is the total number of comparisons (12 comparisons), and *p_i_’* is the p-value adjusted for the number of the multiple hypothesis tests. We use a family wise error rate of 0.05 to reject the hypothesis.

In addition to using one-sided tests on accuracy and specificity, we tested whether each program had zero bias or equivalently, whether the mean error terms equals zero via a two-tailed t-test (Fig. 3A, C, E, G, Bonferroni adjustment applied for 4 comparisons, one for each program). For illustrative purposes, the accuracy data from Fig. 3 was also visualized as a series of Bland-Altman plots^26^ (Supplemental Fig. 4), where the difference between a program’s output metric and ground-truth was plotted against their mean value for each image (meaning Y*_i,j_-G_i,j_* was plotted against (Y*_i,j_*+*G_i,j_)/2*). This analysis offers an illustration of a method commonly used in science to compare measurement methods, and highlights the difficulty in interpreting results from several measurement methods, each with a collection of output variables.

The accuracy of vessel structure metrics was compared before and after user curation with paired t-tests with a Bonferroni adjustment to the p-value for 4 comparisons, one for each of the vessel structure metrics. Since REAVER’s automated results were extremely accurate (Fig. 3) compared to the other programs, and consequently the potential effect size for improvement from user curation was small, the analysis was conducted with default internal image processing parameters with REAVER and then repeated with a separate set of substandard parameters: comparing *A^B,D^_i,r_* to *A^F,D^_i,r_* (Fig. 5A-D) and then separately comparing *A^B,S^_i,r_* to *A^F,S^_i,r_* (Fig. 5E-H).

All of the test statistics examined may not follow a normal distribution. Nevertheless, the sample size of 36 images ensures the robustness of the paired t-test to the violation of the normality assumptions because of the central limit theorem^36^.

Images of vessel architecture in the retina was analyzed across distinct spatial locations with regards to radius and depth from the optic nerve (Supplemental Fig 3) and processed with default REAVER parameters. Each of the vessel structure metrics (vessel length density, vessel area fraction, branchpoint count, vessel radius, and others developed previously^2^) were compared between the six locations in the tissue (inner radial region, superficial depth; inner radial region, intermediate depth; inner radial region, deep depth; outer radial region, superficial depth; outer radial region, intermediate depth; outer radial region, deep depth) with pairwise 2-sample t-tests using a Bonferroni correction as stated in Equation 11 (15 comparisons between 6 locations). A principle components analysis was conducted to visualize the qualitative separation of groups across dimensions that maximizes separation^37^.

### Ranking Algorithm

Test results were ranked using a simplified version of an algorithm developed previously^34^, using a conservative approach that assigns the same rank for instances of rank ambiguity. Ranks are assigned by converting the statistical conclusions of pairwise one-sided tests into a dominance matrix, denoted by ***D*,** where *D_i,j_* is set to 1 if we reject the null hypothesis that program *i* is no less than program *j*, and set to zero otherwise, with zeros also found along the principal diagonal.

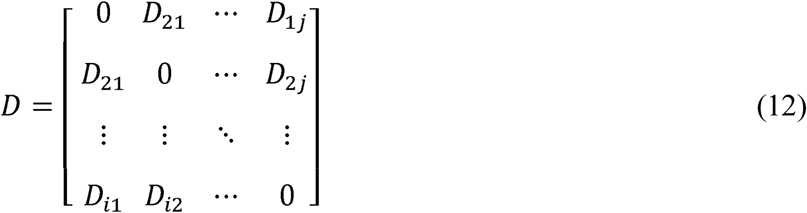

The wins for each program are summed across columns into a column vector *w* of wins, and then values are sorted from most to least as *w’.*

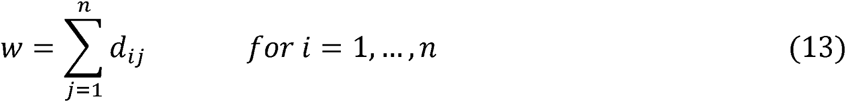

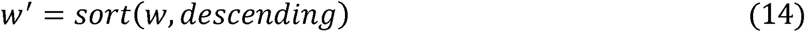

Standard competition ranking (“1224”)^38–41^ is then applied to *w’* to yield *r’,* an ordered vector of assigned ranks, with weak ranking allowing ties to be assigned with the same ranks. As such, each item’s rank is 1 plus the number of items ranked above it.

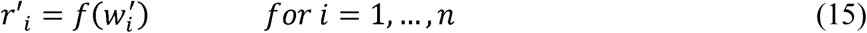

This ranking function is defined as *f* : *w’* → *r’*, and applied to each element of *w’* according to the following formula:

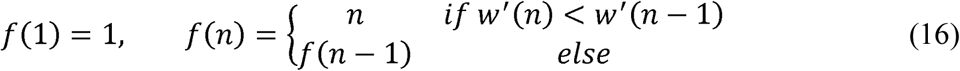

We define an inverse function of the sorting function in equation 14, that restores the ordered vector of assigned ranks from *r’* to yield *r*, a vector of ranks with the original order of programs (found in *w*):

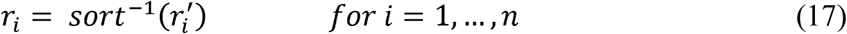

Programs with the top rank are annotated with an up triangle in the plots of vessel structure metric accuracy and precision, segmentation accuracy, specificity, and sensitivity, along with execution time. We note that there are cases when programs assigned the second-best rank, yet are not significantly different from the highest ranked program. It could be argued that the second ranked program should also be considered a top performer under these circumstances, and should be examined on a case-by-case basis based on the effect size and variance.

## Author Contributions Statement

B.A.C and R.D. developed software, analyzed data, and wrote manuscript. C.M. generated data and aided with manuscript. T.Z. aided with statistical analysis and interpretation. P.Y. gave input on drafting of manuscript. S.P.C supervised the project.

## Acknowledgements

We thank Dr. Gustavo Rohde of University Virginia for providing key feedback for improvements to the manuscript.

