## Supplementary material for "REAVER: Improved Analysis of High-resolution Vascular Network Images Revealed Through Round-robin Rankings of Accuracy and Precision": Online Supplement

**Short Running Lead**: REAVER: Image Analysis of Vessel Architecture

**This supplement contains:**

Supplementary Figures 1-5

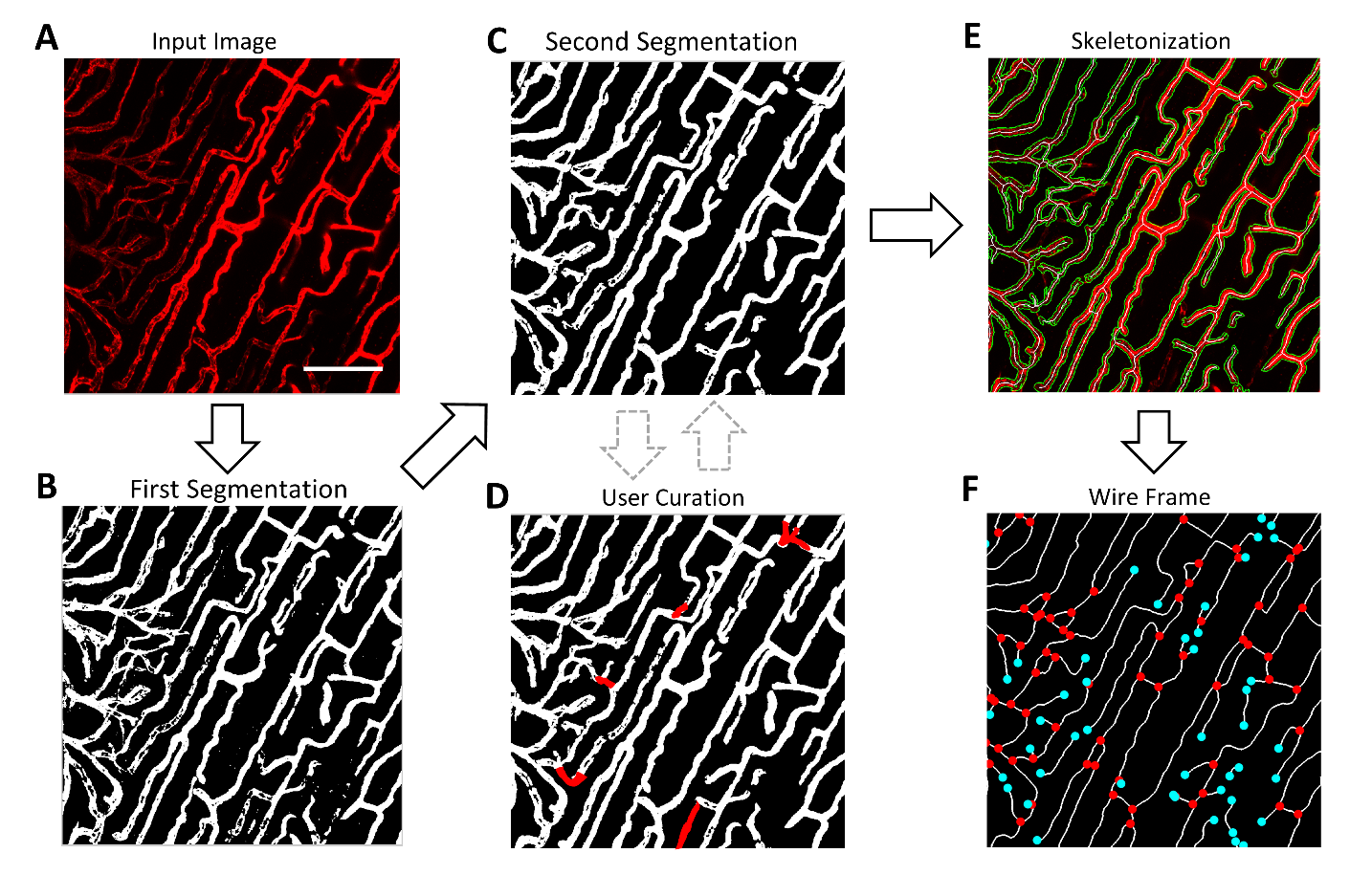

**Supplementary Figure 1: flowchart of REAVER’s algorithm.** Graphical representation of the basic steps in REAVER’s image analysis algorithm, demonstrated with an (**A**) example image analyzed with (**B**) primary segmentation, (**C**) secondary segmentation, (**D**) optional user curation, (**E**) skeletonization, and (**F**) analysis of skeleton architecture.

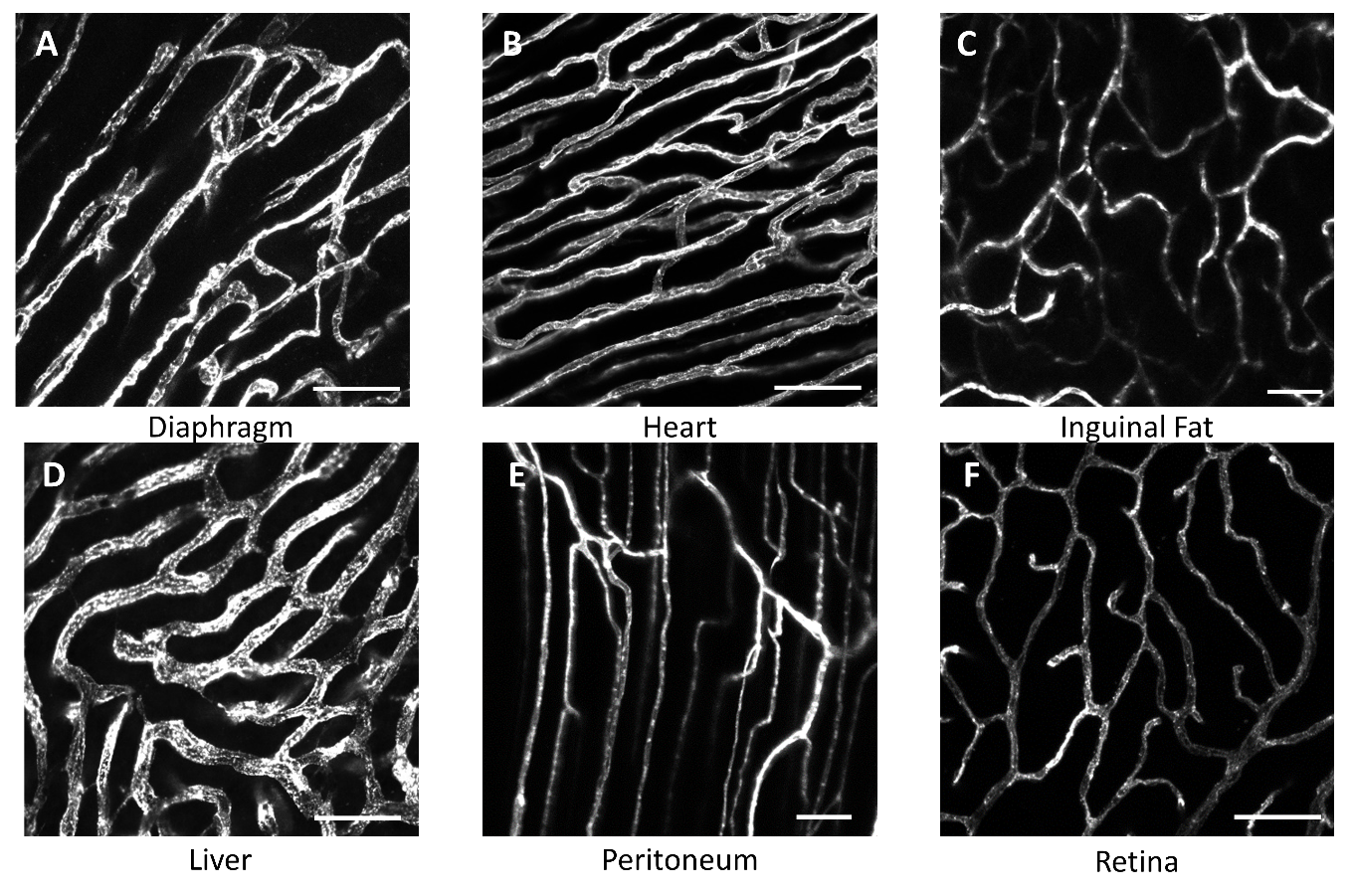

**Supplementary Figure 2: Representative images from tissue used in benchmark dataset.** Six images were acquired from (**A**) diaphragm, (**B**) heart, (**C**) inguinal far, (**D**) liver, (**E**) peritoneum, and (**F**) retina with a mixture of 20x and 60x fields of view (scale bar 25 um).

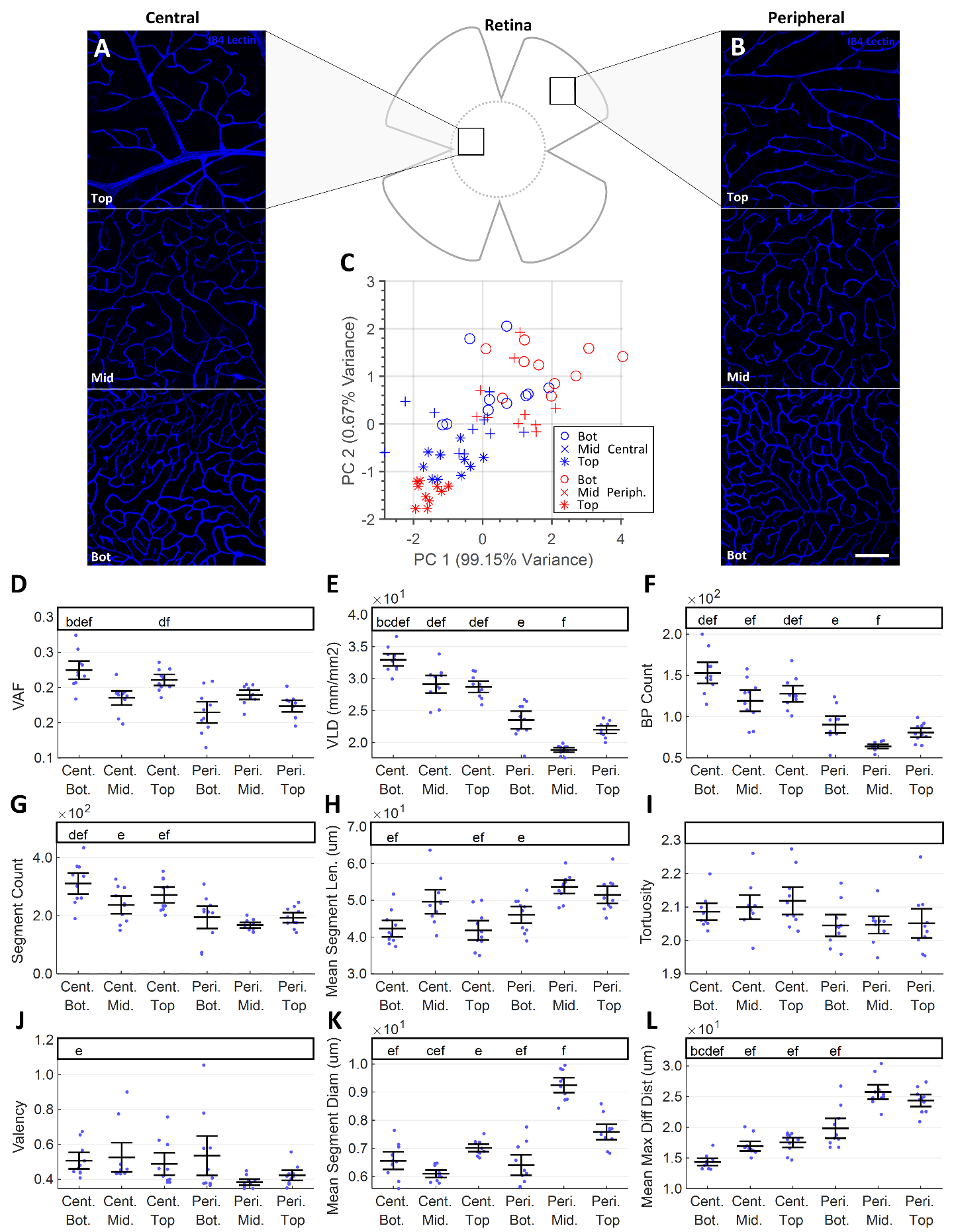

**Supplementary Figure 3: REAVER reveals spatial heterogeneity of vascular network between distinct locations in mouse retina**. A dataset of 30 images were acquired from a lectin perfused mouse retina either (**A**) close to the optic disk, or (**B**) radially towards the periphery of the retina, encompassing each of the three vascular layers (superficial, intermediate, and deep plexus). (**C**) Principle components analysis of vessel architecture metrics leads to partial separation of images acquired from distinct physiologic locations based on a set of metrics. Metrics quantified include (**D**) vessel area fraction, (**E**) vessel length density, (**F**) branchpoint count, (**G**) segment count, (**H**) tortuosity, (**I**) valency, (**J**) vessel diameter, and (**K**) max diffusion distance (pairwise paired 2-tail t-tests, with 15 comparison Bonferroni correction, α=0.05, N=10).

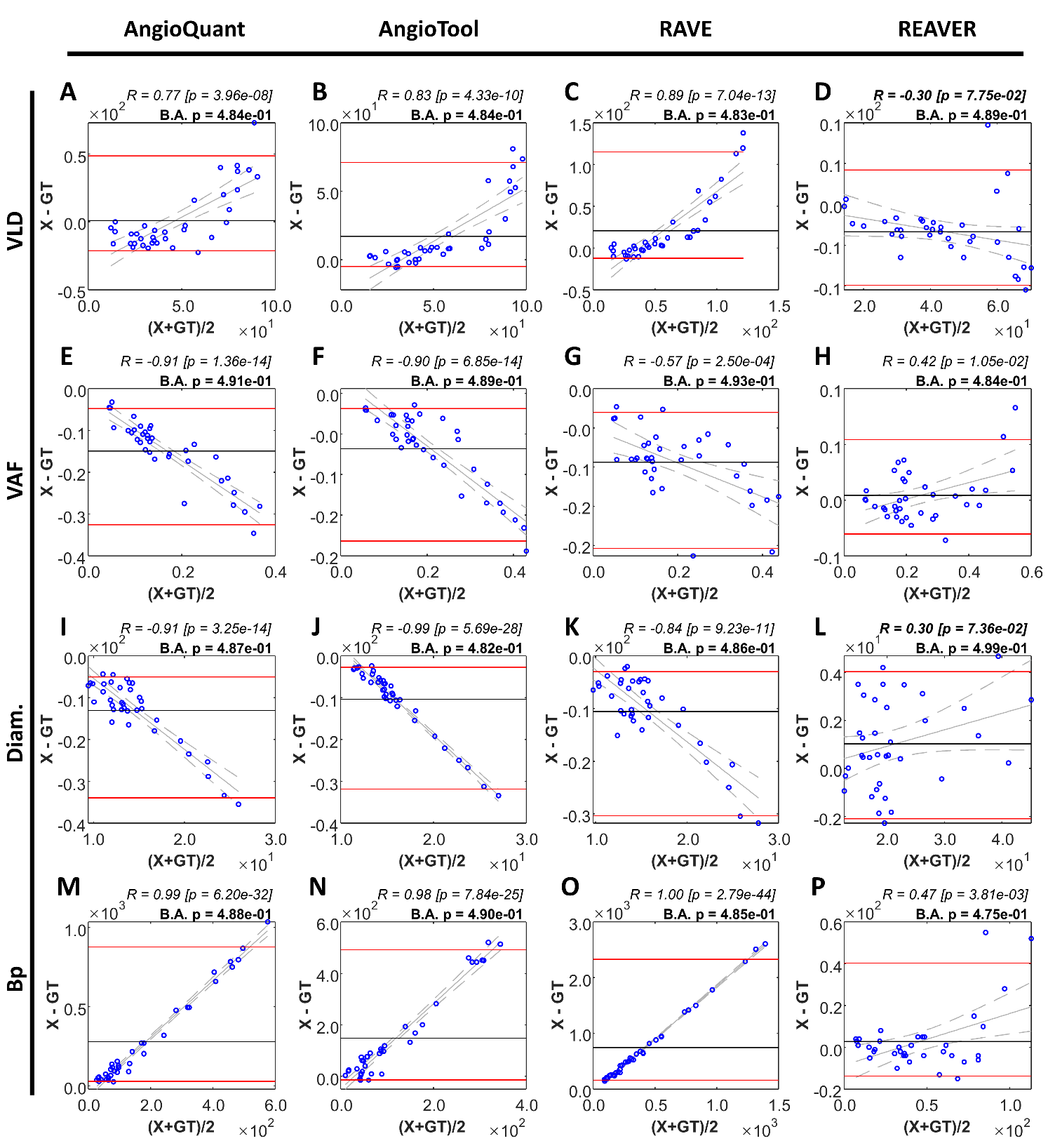

**Supplementary Figure 4: Bland-Altman plots of each metric and program compared to manual analysis**. Bland-Altman plots comparing automated results from 4 vascular image processing pipelines compared to manual analysis as a gold standard, including (**A-D**) vessel length density, (**E-H**) vessel area fraction, (**I-L**) vessel diameter, and (**M-P**) branchpoint count. Black horizontal line is the mean difference between measurements, red lines demarcate 95% agreement interval calculated with bootstrapping (1e6 samples), grey line is best fit of linear model to data, with dashed grey lines the 95% confidence interval of the fit.

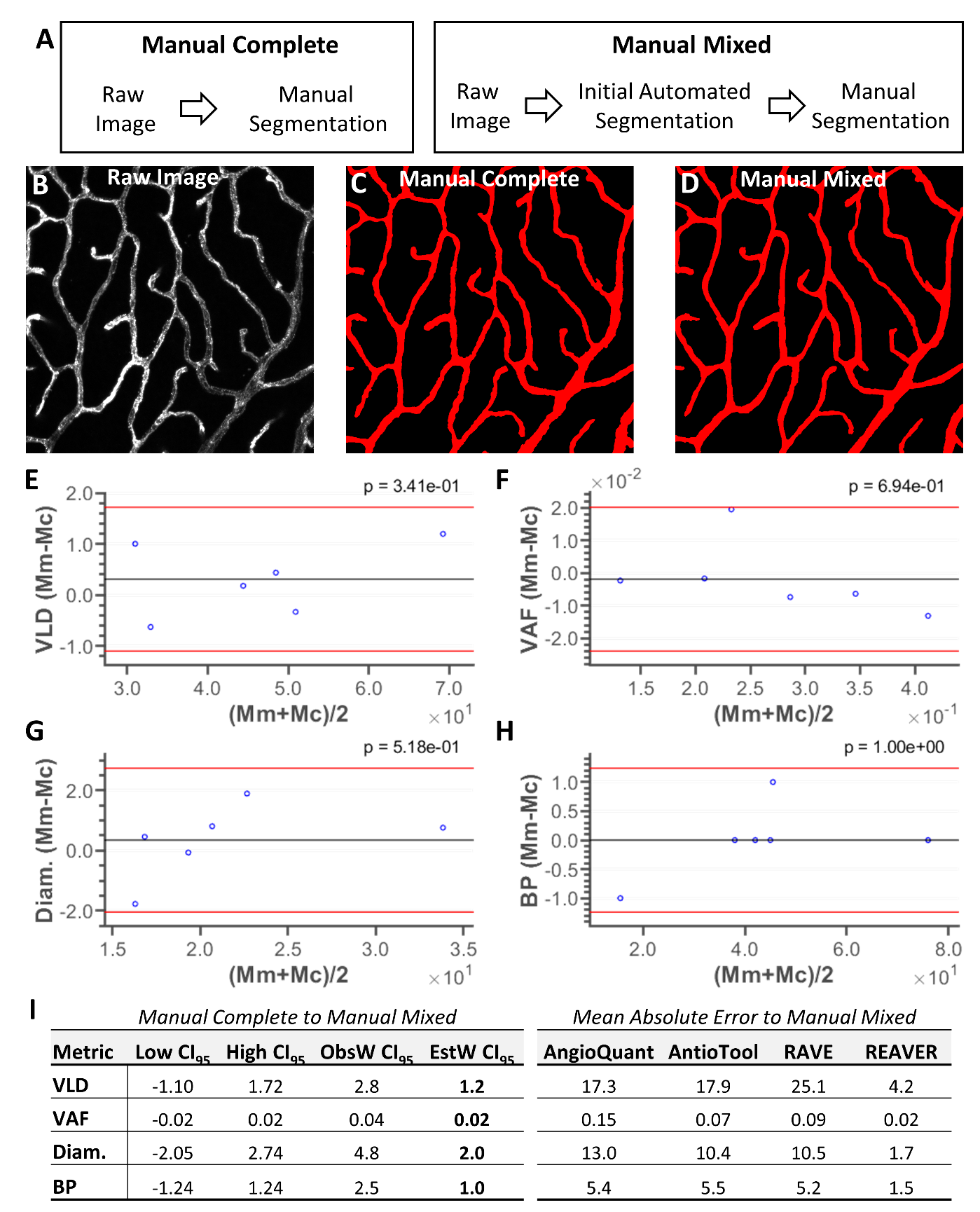

**Supplementary Figure 5: Disagreement between ground-truth segmentation methods is minor compared to effect sizes between programs**. (**A)** Completely manual annotation of vessel segmentation (Mc) was compared to mixed manual (Mm, the method used for all images in the image benchmark dataset) to check if bias with the mixed manual method could have unfairly favored REAVER. (**B-D**) Example input image with segmentations from both methods. (**E-H**) Bland-Altman plots examining disagreement between manual mixed and manual complete segmentation methods with upper and low limits of 95% confidence interval (red) for each of the vessel architecture metrics (p-values displayed from two-sided 1-sample t-tests, N=6 images, one from each tissue type, no multiple comparisons corrections to represent a conservative interpretation). (**I**) Table of the confidence intervals and widths of the disagreement between ground-truth analysis methods (ObsW CI_95_, observed width 95% confidence interval, normality assumed, N=6 images, one from each tissue type, no multiple comparisons corrections to represent a conservative interpretation). The confidence intervals of disagreement between manual analysis methods for the entire benchmark dataset were estimated based on increasing the sample size from 6 to 36 images (decreasing the interval by 1/√n) with sample standard deviation fixed (EstW CI_95_, estimated width 95% confidence interval,). Mean absolute error observed in each of the automated programs to mixed manual ground-truth displayed to allow comparison to confidence intervals to effect sizes (N=36 images, data reproduced from Fig. 3).
